# A dynamical system model for predicting gene expression from the epigenome

**DOI:** 10.1101/2020.08.03.234740

**Authors:** James D. Brunner, Jacob Kim, Timothy Downing, Eric Mjolsness, Kord M. Kober

## Abstract

Gene regulation is an important fundamental biological process. The regulation of gene expression is managed through a variety of methods including epigentic processes (e.g., DNA methylation). Understanding the role of epigenetic changes in gene expression is a fundamental question of molecular biology. Predictions of gene expression values from epigenetic data have tremendous research and clinical potential. Despite active research, studies to date have focused on using statistical models to predict gene expression from methylation data. In contrast, dynamical systems can be used to generate a model to predict gene expression using epigenetic data and a gene regulatory network (GRN) which can also serve as a mechanistic hypothesis. Here we present a novel stochastic dynamical systems model that predicts gene expression levels from methylation data of genes in a given GRN. Software for dataset preparation, model parameter fitting and prediction generation, and reporting are available at https://github.com/kordk/stoch_epi_lib.

## Introduction

Gene regulation is an important fundamental biological process. [1] It involves a number of complex sub-processes that are essential for development and adaptation to the environment (e.g., cell differentiation [2] and response to trauma [3]). Understanding gene expression patterns has broad scientific [4] and clinical [5] potential, including providing insight into mechanisms of regulatory control [1] (e.g., gene regulatory networks) and a patient’s response to disease (e.g., HIV infection [6]) or treatment (e.g., chemotherapy-induced neuropathic pain [7]). The regulation of gene expression is managed through a variety of methods, including transcription, post-transcriptional modifications, and epigenetic processes [8]. One epigenetic process, DNA methylation, [9] occurs primarily at the cytosine base of the molecule that is adjacent to guanine (i.e., CpG site). While evidence exists to support a relationship between methylation and gene expression, the patterns of these associations can vary. [10] DNA methylation of promoter and gene body regions can act to regulate gene expression by repressing [11]) or activating [12] transcription. For example, higher gene expression can be associated with both decreased [13] and increased [14] methylation in regulatory regions, and with decreased methylation within the gene. [15] These associations vary with the distance from the promoter, [16] as well as between individuals and across tissues. [17]

Predicting gene expression levels from epigenetic data is an active area of research. Recent studies have developed models to predict gene expression levels with deep convolutional neural networks from genome sequence data [18] and with regression models from methylation data. [19, 20] Earlier studies developed models to predict expression status (e.g., on/off or high/low) with gradient boosting classifiers from histone modification data [21], with machine learning classification methods from methylation data [22], and from methylation and histone data combined. [23] However, these studies exclusively use a statistical approach to predicting gene expression. One limitation of the deep learning approaches is in the interpretation of the results. [24] A limitation of the linear model approaches are their inability to provide information regarding promoter binding events and the regulatory activities of the system. Neither of these approaches can provide a biological model to explain the expression estimates.

To address these limitations, we developed a dynamic interaction network model [25] that depends on epigenetic changes in a gene regulatory network (GRN). Dynamical systems integrate a set of simple interactions (i.e., transcription factor (TF) binding to a promoter region and subsequent gene expression) across time to produce a temporal simulation of a physical process (i.e., gene regulation in a given GRN). Therefore, the predictions of a dynamical systems model (e.g., TF binding and unbinding events, gene expression levels) emerge from a mechanistic understanding of a process rather than the associations between data (e.g., predicting an outcome from a set of predictor variables). A dynamical systems model can predict gene expression using epigenetic data and a GRN by simulating hypothesized mechanisms of transcriptional regulation. Such models provide predictions based directly on these biological hypotheses, and provide easy to interpret mechanistic explanations for their predictions. The dynamical systems approach offers a number of unique characteristics. First, a stochastic dynamical system provides us with a distribution of gene expression estimates, representing the possibilities that may occur within the cell. Next, the mechanistic nature of the approach means that the model can provide a biological explanation of its predictions in the form of a predicted activity level of various gene-gene regulatory interactions. Finally, a dynamical systems approach allows for the prediction of the effects of a change to the network. To our knowledge, there are no studies that have taken a dynamical systems approach to predicting gene expression from methylation data and a GRN.

Given the opportunity presented by dynamical systems approaches and the potential practical utility, we present a novel stochastic dynamical systems model for predicting gene expression levels from epigenetic data for a given GRN, along with a software package for model parameter fitting and prediction generation (available at https://github.com/kordk/stoch_epi_lib).

## Methods

Here we use a dynamical systems approach to develop and fit a model to predict gene expression levels and transcription factor binding affinities from methylation data Fig. 1.

**Fig 1.**
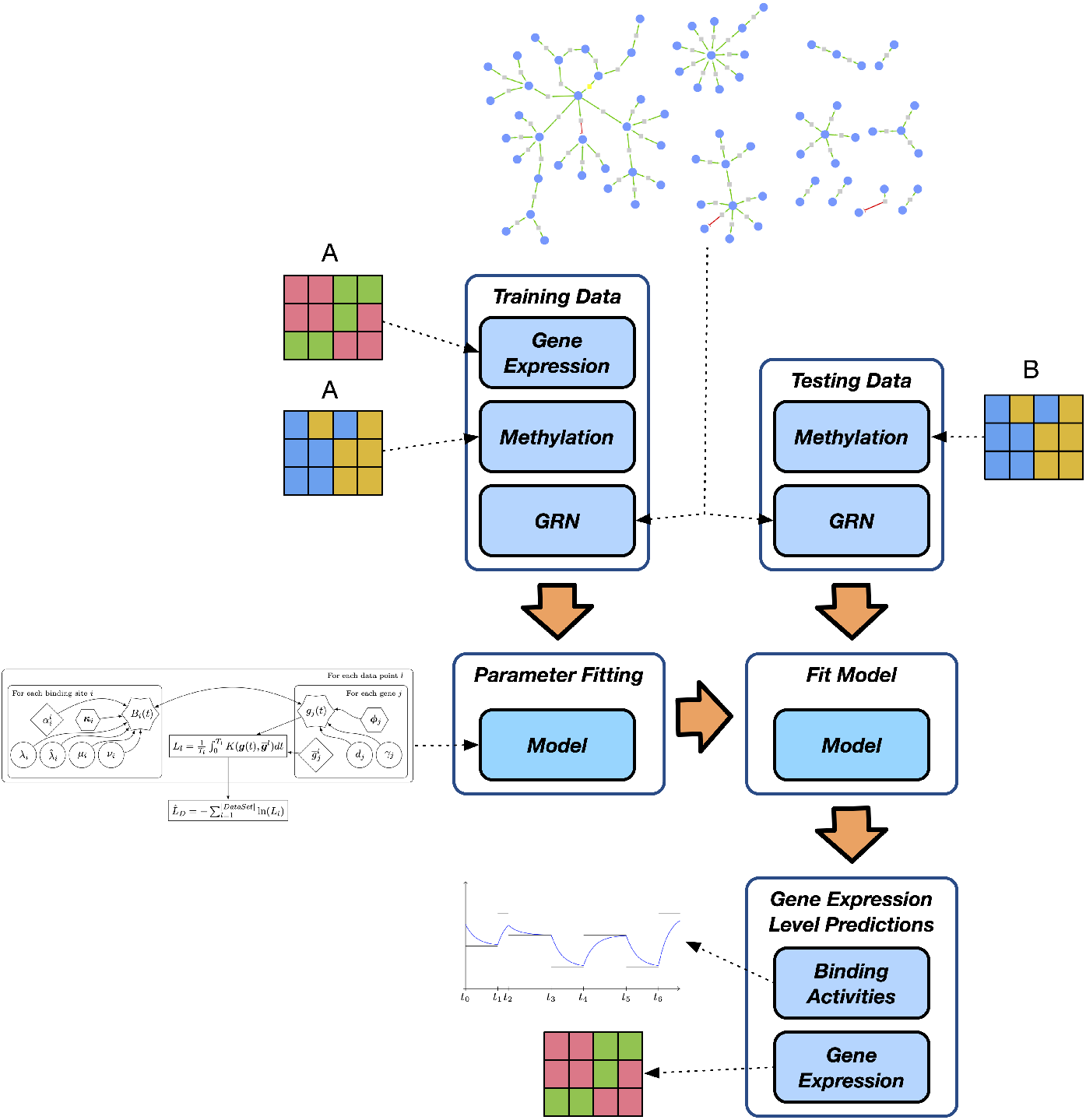
An overview of our approach using a dynamical systems model to predict gene expression using a gene regulatory network and methylation data. Gene expression and methylation data from training set A is used to fit the parameters of the model. Gene expression and binding activities are predicted using the fit model and methylation data from testing set B.

### Model Equations

We model gene regulation using a piecewise-deterministic Markov process (PDMP) as introduced in Davis 1984 [26] (see also [27, 28]) given by the equations:

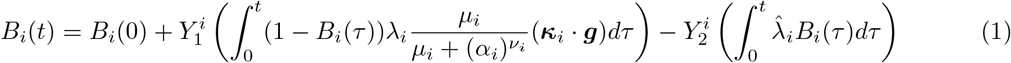

and

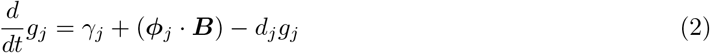

where *B*_*i*_(*t*) ∈ {0, 1}, is a boolean random variable representing the bound/unbound state of a binding site region of DNA and *g*_*i*_ is the transcript amount the genes modeled. Equation (1) is given as the sum of two Poisson jump processes 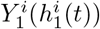 and 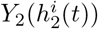 which take values in ℤ_≥0_, and are piecewise constant between randomly spaced discrete time points (the binding and unbinding events) [29].

The propensities 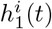 and 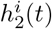 are taken to be linear functions of the available transcription factors, which is assumed to be the same as the transcript variables *g*_*j*_. We take the values *κ*_*ij*_ ∈ {0, 1}; these parameters along with the set of *ϕ*_*ji*_ represent the structure of the underlying gene regulatory network.

We include the term

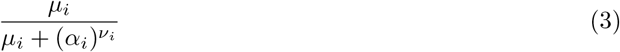

to represent the impact of epigenetic modification on the binding propensity of transcription factors. In this term, *α*_*i*_ is the measured epigenetic modification to a transcription factor binding site (e.g. percentage of methylated bases). Equation (3) is a sigmoidal function which is either strictly increasing or strictly decreasing depending on the sign of *ν*_*i*_.

Finally, we use a linear ODE for the value of the transcripts *g*_*j*_. We take *ϕ*_*ji*_ ∈ {−1, 0, 1} based on the structure of the underlying gene regulatory network. We include baseline transcription *γ*_*j*_ and decay *d*_*j*_. Because we use a linear ODE in Eq. (2), we can solve exactly between jumps of ***B***.

It is common practice in the study of reaction networks modeled as stochastic jump processes to represent the process using so called “chemical master equation” [29, 30] (i.e. the Kolmogorov forward equation for the jump process), which can be used within optimization methods to learn parameters for the system [31]. The generator for a PDMP can be defined (see Azaïs 2014 [32] for details). We define a density *P* (*B*^*i*^, ***g***(*t*), *t*) = *P*^*i*^(***g***), *i* = 1, *...,* |***B***| for each possible state ***B***^*i*^ of ***B*** such that 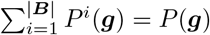 is the probability distribution for the vector ***g***, and each *P*_*i*_ satisfies

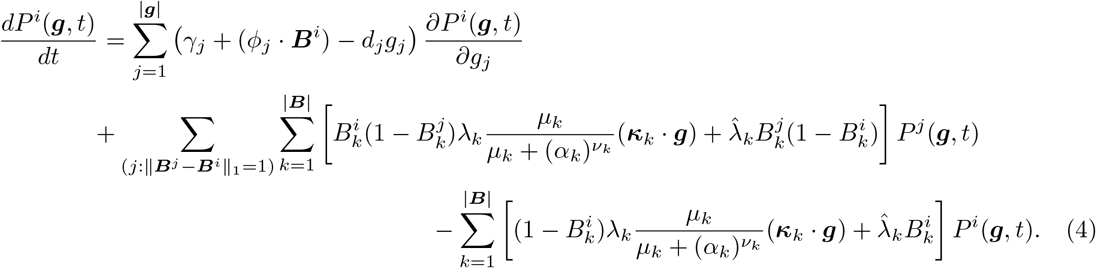

### Model Parameter Estimation

The parameters *κ*_*ij*_, *ϕ*_*ji*_ and *γ*_*j*_ are determined by the structure of the underlying gene regulatory network and the epigenetic parameter *α*_*i*_ is assumed measurable. We estimate the parameters *λ*_*i*_, 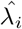, *μ*_*i*_, *ν*_*i*_ and *d*_*j*_. We estimate these parameters using a negative log-likelihood minimization procedure using stochastic gradient descent. Sample paths used to estimate the gradient of the likelihood are generated using one of two modified forms of Gillespies stochastic simulation algorithm (SSA) [33] which handle time-dependent jump propensities by adding an ODE to the system [27, 30] or by rejecting jumps chosen as in the standard SSA [34]. This procedure involves approximating the gradient of the map from parameter set to log-likelihood so that we may use a gradient descent method.

We can compute a log-likelihood for a set of paired epigenetic and transcription samples by time averaging a sample path against a Gaussian kernel. We estimate the likelihood of a sample of transcript data 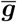 of *d* genes using a model realization ***B***(*t*), ***g***_***θ,α***_(*t*) as follows:

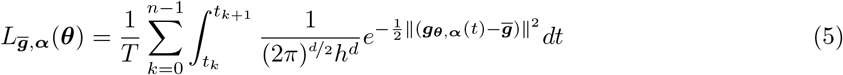

where *h* is some smoothing parameter, ***θ*** is an *n*-dimensional vector of model parameters where *n* is the sum of the sizes of sets of *λ*_*i*_, 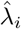, *μ*_*i*_, *ν*_*i*_, and ***α*** represents the epigenetic parameters *α*_*i*_ used by the model. For a dataset *D* consisting of *n* sets of matched pairs of transcription and epegenetic data 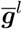, ***α***^*l*^, we define the negative log-likelihood as:

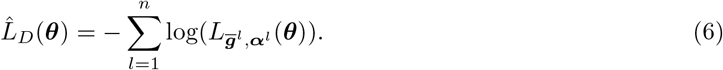

We note that 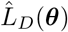 depends on computing ***g***_***θ,α***_(*t*) from a realization of the stochastic process, and so depends on a set ***ω*** of random numbers. So that we are minimizing a deterministic map, we choose ***ω*** once and use this same random vector to generate every realization needed in computing the estimate 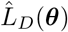. We make this explicit with the notation *L*_*D*,***ω***_(***θ***).

In Fig. 2, we give a schematic representation of how 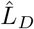 is estimated from a set of realizations of the model, each realization corresponding to a single data sample. Details of the gradient estimation are given in supplemental file S1.

**Fig 2.**
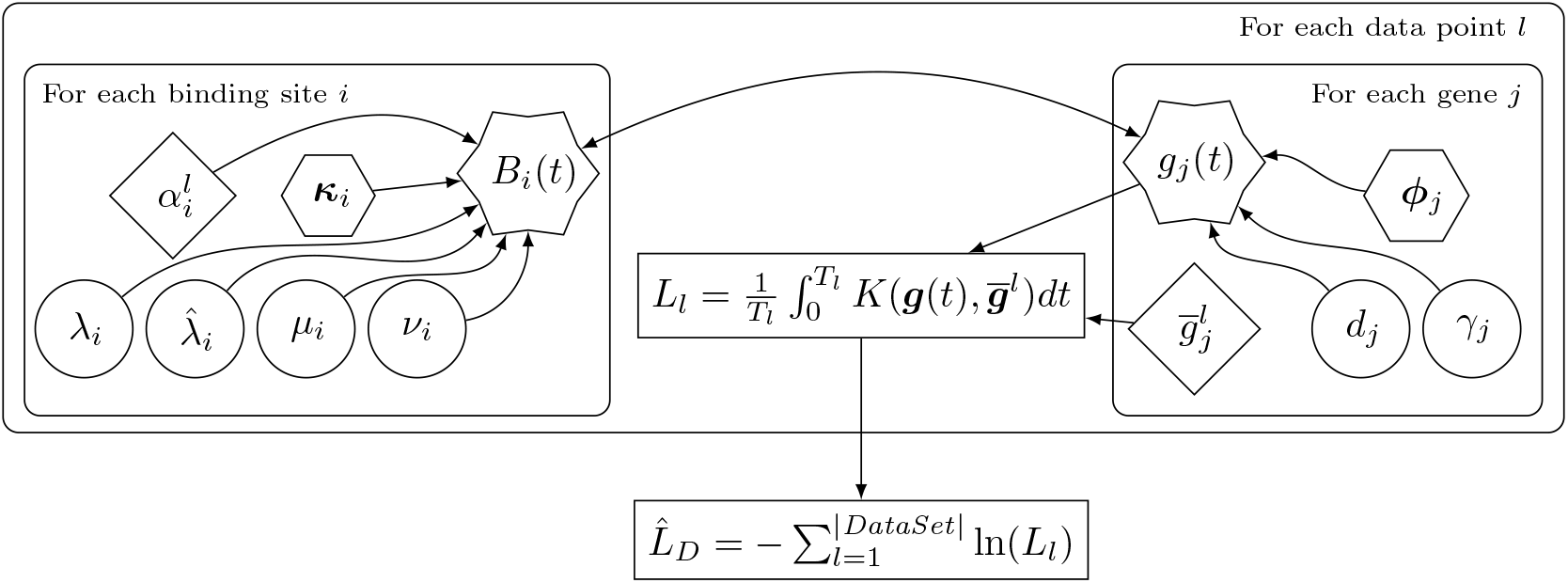
Plate diagram of the process to estimate total likelihood of a data set according to our model. Parameters in diamonds are read from data, parameters in hexagons are determined by the structure of the network, parameters in circles must be fit to the model by maximizing likelihood over a training data set, and parameters in stars are the state variables of the dynamical model. Notice that the dynamical model implies that the stat variables depend on each other, meaning this network of dependence is *not acyclic*. The kernel *K*(*x, y*) used to estimate likelihood is Gaussian.

### Evaluation

#### Gene Regulatory Network

Gene to gene interactions were defined using the Discriminant Regulon Expression Analysis (DoRothEA) framework. [35] Transcription factor (TF) to target interactions were identified as those with the DoRothEA highest confidence interaction classification and scored as 1 or −1 for upregulating and downregulating, respectively. Binding site to target edges (*ϕ*) were defined by CpG methylation sites which were associated with changes in transcript expression (eCpG). [36]

#### Dataset

Matched epigenetic and gene expression data were obtained from whole blood from participants in the Grady Trauma Project (GTP) study (n=243 participants). Methylation data were obtained from the NCBI Gene Expression Omnibus (GEO) (GSE72680) and measured using the HumanMethylation450 BeadChip (Illumina, San Diego, CA). Methylation status was quantified as a beta score. A total of 19,258 eCpG probes were identified. Beta scores for CpG sites within the same region for a gene (i.e., classified as either ‘Promoter’ or ‘TSS’ [36]) were aggregated together as the mean. Gene regions where no DNA methylation data were collected were excluded. A total of 1,885 regions were identified.

Gene expression data were obtained from GEO (GSE58137) measured with the HumanHT-12 expression beadchip V3.0 (Illumina, San Diego, CA). Intensity scores were log2-transformed (mean expression intensity = 189.96, IQR = 49.88 to 106.60). Gene expression probes were first annotated to ENTREZ ID and then annotated to the symbol using the HUGO database. [37]

For evaluation, we identified a set of genes previously identified as deferentially expressed in individuals with PTSD as compared to controls (n=524). [38] Of these, we identified 278 TF to target mappings using the DoRothEA framework. We then used this list of genes to identify additional targets to include beyond initial list. The final set included 252 TF to target relationships comprised of 303 unique target genes. A GRN was built using these 303 genes as input producing a final network with 74 genes with 65 sites (Fig. 3). Of these 74 genes, 29 had sufficient data and regulatory information (i.e., methylation and gene expression data for all individuals, an eCpG binding site, and a TF to gene relationship) for which parameters could be estimated and expression distributions generated.

**Fig 3.**
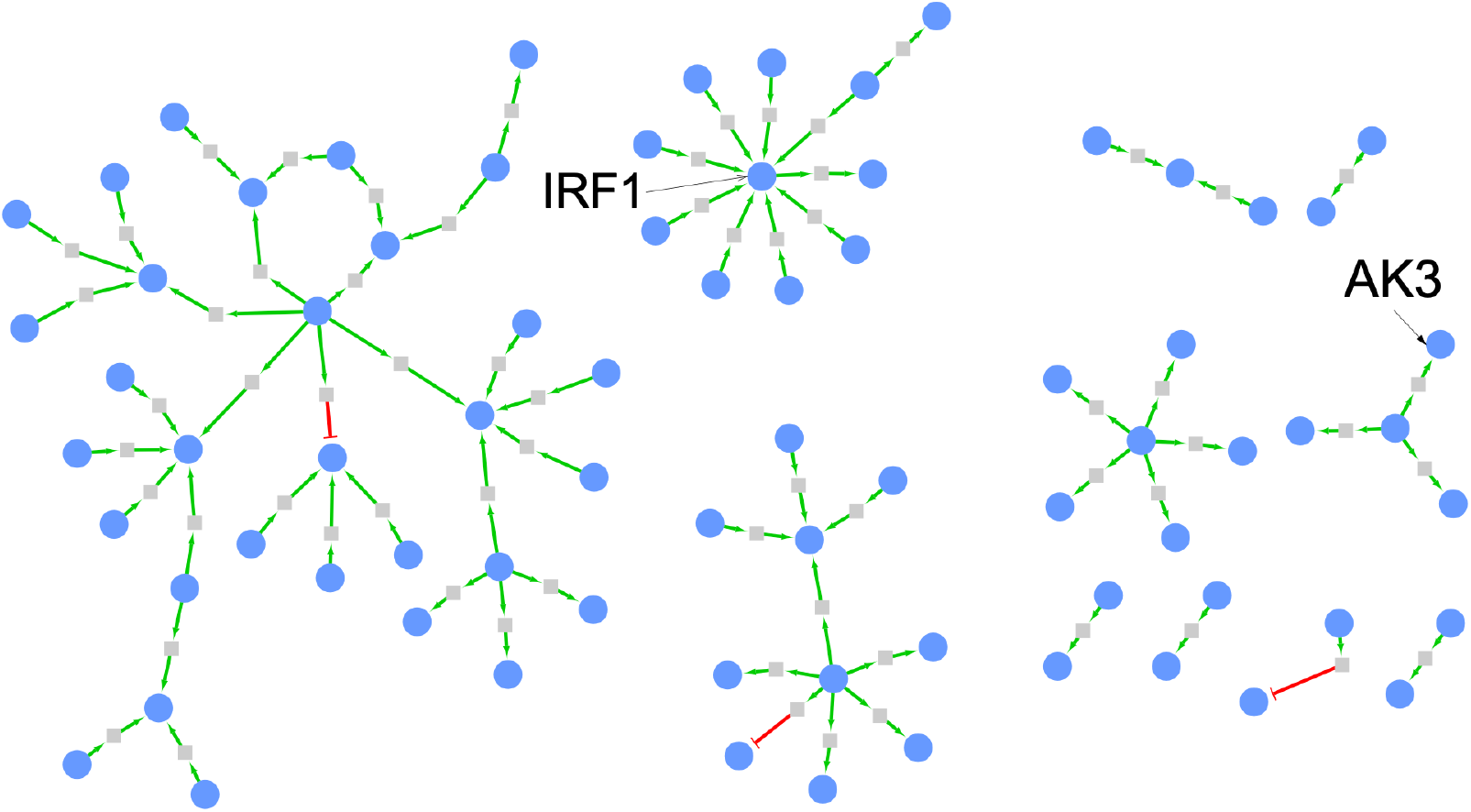
Bipartite network corresponding to the gene regulatory network based on differential expression in individuals with PTSD. This network contains seventy-four genes and sixty-five sites. Of these, twenty-nine genes had sufficient regulatory information (i.e., an associated binding site and transcription factor) for which parameters could be estimated and expression distributions generated. Blue circles are genes, grey boxes are binding sites. Green arrows are activating and red ’T’s are inhibitory. Black arrows point to AK3 and IRF1.

#### Cross Validation

Matched gene expression and methylation data from participants measured for expression (n=243) were used for evaluation. This primary dataset was split into training and testing datasets, containing 80%/20% (n=195, n=48 samples, respectively). To avoid the impact of a particular split, we repeated the shuffle process 100 times. [39] For each split of the data, parameter estimation was performed on the training set and equilibrium distributions of the predicted expression levels were generated using the testing set. For every round of cross-validation, the error in prediction was evaluated as the root mean square error (RMSE) [40] between the observed and the estimated expression from our model. To rank methods, the RMSEs (mRMSE) was averaged for each method across the 100 shuffles.

#### Model Comparison

To evaluate the performance of our gene expression predictions we generated linear regression models using the *scikit-learn* software package for python [41]. Based on previous studies that developed prediction models for gene expression using methylation data, [19, 20] we generated prediction models using LASSO, Multi-task LASSO, Elastic Net, and Multi-Task Elastic Net. The structural parameters for these models (i.e. penalty parameter and *l*_1_-ratio parameter) were determined using scikit-learn’s cross-validation methods with the entire data set. Finally, we fit a null model that is the average of the expression values from the training set. It is the prediction of expression values without any other variables in the model. Models were generated for each of the 100 data train/test shuffles used in our fitted model.

To evaluate the performance of our fitting procedure on gene expression predictions we generated predictions using a randomly generated parameter set a for each of the previously generated splits. Ten random estimates were generated for each shuffle giving 1000 predictions for each gene generated using random parameters. Parameters were estimated for all genes using the procedure detailed in Supplemental File 1.

## Results

Across the final models, our fitted parameter model performed the best (Table 1, Fig. 4). Across all 29 genes, our model outperformed the null model as well as the four linear regression models Fig. 5. On average, our model outperformed the best performing linear regression model (i.e., ElasticNet) by a factor of 2. 16 after parameter fitting. The average root mean square errors for each gene across the 100 shuffles is reported in Table 2. We observed the highest performance for AK3 (average RSME = 1.101) and lowest for SCP2 (average RSME = 2.697). In this evaluation, our model biases towards underestimating the expression levels (see supplemental file).

**Table 1.** Summary of Average mRMSE of 100 splits of training and testing data across 29 genes for seven prediction models (i.e., our fitted model, multitask (MT) elastic net, elastic net, multitask LASSO, LASSO, our model with random parameters, and training data transcript average value (null)).

|  | Model | MT ElasticNet | ElasticNet | MT LASSO | LASSO | Null (Mean) | Random |
| --- | --- | --- | --- | --- | --- | --- | --- |
| count | 29.000 | 29.000 | 29.000 | 29.000 | 29.000 | 29.000 | 29.000 |
| mean | 2.059 | 4.479 | 4.477 | 4.485 | 4.474 | 4.473 | 8.750 |
| std | 0.571 | 0.898 | 0.879 | 0.899 | 0.870 | 0.897 | 1.444 |
| min | 1.101 | 3.354 | 3.361 | 3.359 | 3.364 | 3.349 | 6.114 |
| 25% | 1.657 | 3.739 | 3.738 | 3.744 | 3.739 | 3.733 | 7.631 |
| 50% | 2.060 | 4.254 | 4.254 | 4.259 | 4.255 | 4.248 | 8.767 |
| 75% | 2.564 | 5.162 | 5.180 | 5.168 | 5.186 | 5.156 | 9.658 |
| max | 3.172 | 6.410 | 6.254 | 6.418 | 6.163 | 6.405 | 12.030 |

**Fig 4.**
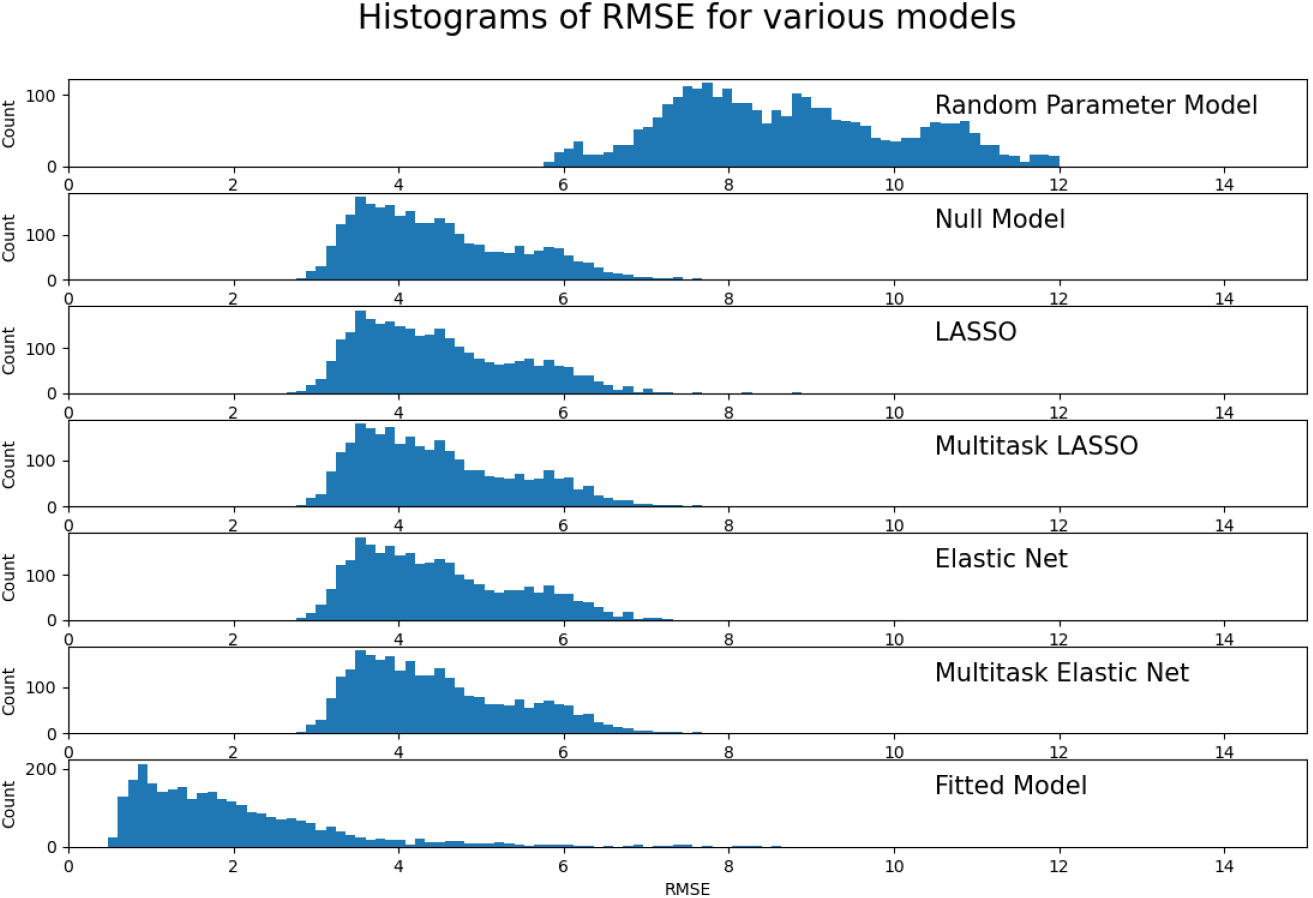
Histogram of all RMSEs across 29 genes and 100 distinct train/test data splits for each model.

**Fig 5.**
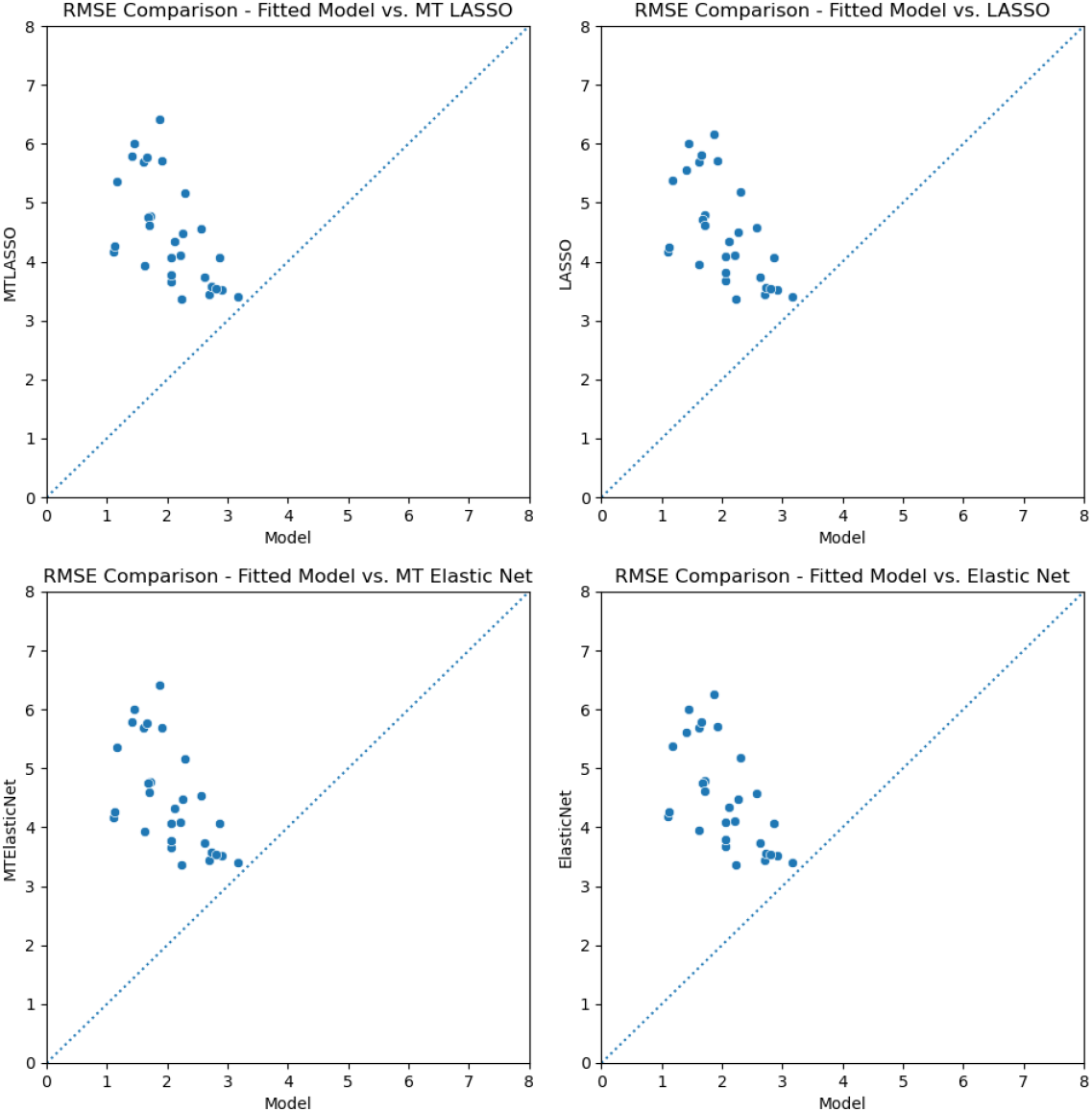
Comparison of average RMSE of our fitted model with four linear regression models for each gene.

**Table 2.** Average root mean square errors for each gene across the 100 shuffles for 7 models - our fitted model, multitask elastic net, elastic net, multitask LASSO, LASSO, training data transcript average value (Null), our model with random parameters.

|  | Model | MT ElasticNet | ElasticNet | MT LASSO | LASSO | Null (Mean) | Random |
| --- | --- | --- | --- | --- | --- | --- | --- |
| LDHA | 1.917 | 5.698 | 5.712 | 5.709 | 5.719 | 5.688 | 10.680 |
| NR1D2 | 2.215 | 4.095 | 4.107 | 4.100 | 4.111 | 4.090 | 7.602 |
| SREBF1 | 2.863 | 4.058 | 4.071 | 4.063 | 4.075 | 4.053 | 6.783 |
| CD4 | 1.711 | 4.768 | 4.786 | 4.776 | 4.792 | 4.762 | 8.794 |
| RRM2B | 2.060 | 3.659 | 3.677 | 3.663 | 3.682 | 3.655 | 8.195 |
| SLC20A1 | 1.614 | 5.687 | 5.688 | 5.697 | 5.691 | 5.679 | 10.787 |
| RPL39L | 2.235 | 3.354 | 3.361 | 3.359 | 3.364 | 3.349 | 8.169 |
| AK3 | 1.101 | 4.166 | 4.179 | 4.174 | 4.167 | 4.159 | 9.617 |
| MT1X | 1.128 | 4.254 | 4.254 | 4.259 | 4.255 | 4.248 | 9.675 |
| ZNF654 | 2.620 | 3.739 | 3.738 | 3.744 | 3.739 | 3.733 | 7.789 |
| ALOX5 | 2.911 | 3.524 | 3.523 | 3.527 | 3.523 | 3.522 | 7.071 |
| CD19 | 1.414 | 5.789 | 5.619 | 5.795 | 5.555 | 5.783 | 9.092 |
| FBXO32 | 1.673 | 4.749 | 4.755 | 4.757 | 4.720 | 4.741 | 8.906 |
| SCP2 | 2.697 | 3.444 | 3.446 | 3.448 | 3.447 | 3.440 | 7.625 |
| CCM2 | 1.620 | 3.932 | 3.948 | 3.936 | 3.953 | 3.929 | 10.344 |
| CTSH | 2.064 | 4.064 | 4.086 | 4.071 | 4.094 | 4.058 | 8.158 |
| FCER1A | 1.446 | 5.995 | 5.994 | 6.003 | 5.998 | 5.988 | 9.658 |
| ICAM4 | 2.732 | 3.570 | 3.567 | 3.573 | 3.567 | 3.566 | 7.746 |
| VWA5A | 2.805 | 3.534 | 3.534 | 3.539 | 3.535 | 3.529 | 7.452 |
| CYP27A1 | 1.172 | 5.352 | 5.374 | 5.359 | 5.385 | 5.345 | 8.767 |
| BAG3 | 2.564 | 4.545 | 4.569 | 4.552 | 4.576 | 4.538 | 7.179 |
| GSTM1 | 1.657 | 5.766 | 5.794 | 5.772 | 5.805 | 5.761 | 10.950 |
| LTA4H | 2.298 | 5.162 | 5.180 | 5.168 | 5.186 | 5.156 | 12.030 |
| SURF6 | 1.708 | 4.603 | 4.610 | 4.611 | 4.613 | 4.596 | 8.937 |
| IRF1 | 2.264 | 4.469 | 4.485 | 4.474 | 4.490 | 4.465 | 10.887 |
| CXCR5 | 2.057 | 3.773 | 3.799 | 3.779 | 3.806 | 3.768 | 7.631 |
| AHR | 3.172 | 3.393 | 3.396 | 3.397 | 3.397 | 3.389 | 6.114 |
| OAS1 | 1.869 | 6.410 | 6.254 | 6.418 | 6.163 | 6.405 | 9.057 |
| BAK1 | 2.123 | 4.330 | 4.337 | 4.337 | 4.341 | 4.324 | 8.042 |

Comparing the model with randomly generated parameters and fitted parameters reveals that our fitting procedure was effective. We see a 4.24-fold improvement in model performance on average after the fitting procedure. In fact, Fig. 4 demonstrates that, with random parameters, our model is unsurprisingly worse than a linear regression, but our fitting procedure returns a model that outperforms linear regression. Examples of the equilibrium distributions generated from the random parameter for the most accurate predicted gene (i.e., AK3) for two individual patients from different shuffles are shown in Fig. 6.

**Fig 6.**
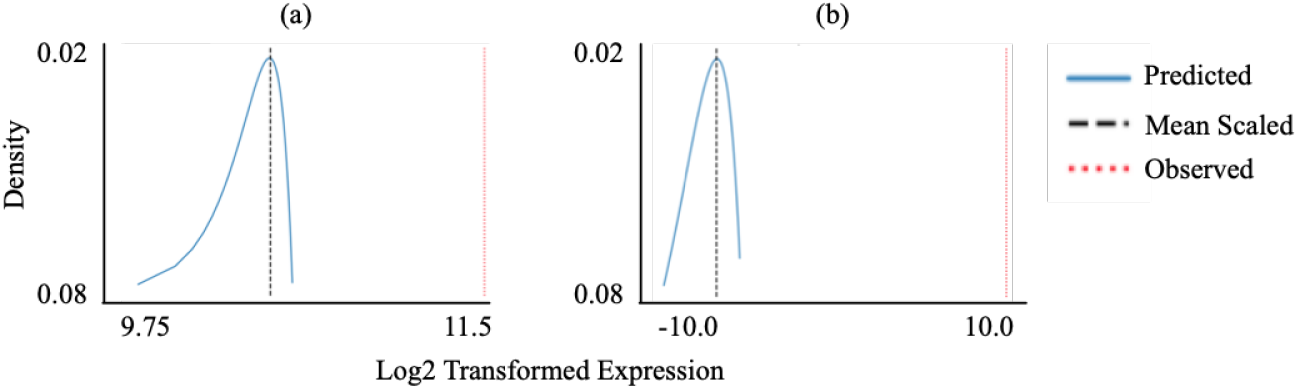
Equilibrium distribution plots generated from random parameters for AK3 for (a) individual ID 6522 for random parameter set 0 in shuffle 0, and (b) individual ID 8331 random parameter set 7 in shuffle 13.

## Discussion

In this study, we demonstrate that gene expression levels can be accurately predicted from methylation state of a promoter region and a GRN. Our model successfully uses quantitative data describing epigenetic modification of transcription factor binding sites to generate a probability distribution which describes the possible level of transcript. To our knowledge, this is the first study to develop and evaluate a stochastic dynamical systems model predicting gene expression levels from epigenetic data for a given GRN.

Overall our model outperforms linear regression approaches in the predictions of the model with fitted parameters (e.g., Fig. 7 a & b) and dramatic improvements to prediction relative to a randomly generated set of parameters (e.g., Fig. 7 c & d). We were able to accurately predict gene expression based on the structure of the GRN which allows for the identification of TF and binding sites that are associated with gene expression levels. For example, our model accurately predicted gene expression levels for both AK3 and IRF1, yet the GRN has different numbers of TF for each (i.e., a single TF for AK3 versus multiple TF for IRF1)(Fig. 3). From our initial list of 302 genes for inquiry, our TF to target and binding site reference data produced a gene regulatory network with 74 genes, of which 29 had sufficient regulatory information to be predicted. Although we were unable to evaluate a more complicated GRN from all reference regulatory data due to computational constraints, we expect that model predictions will improve with additional regulatory information. Future work is needed to improve the computational performance of the implementation to support larger and more complicated GRNs.

**Fig 7.**
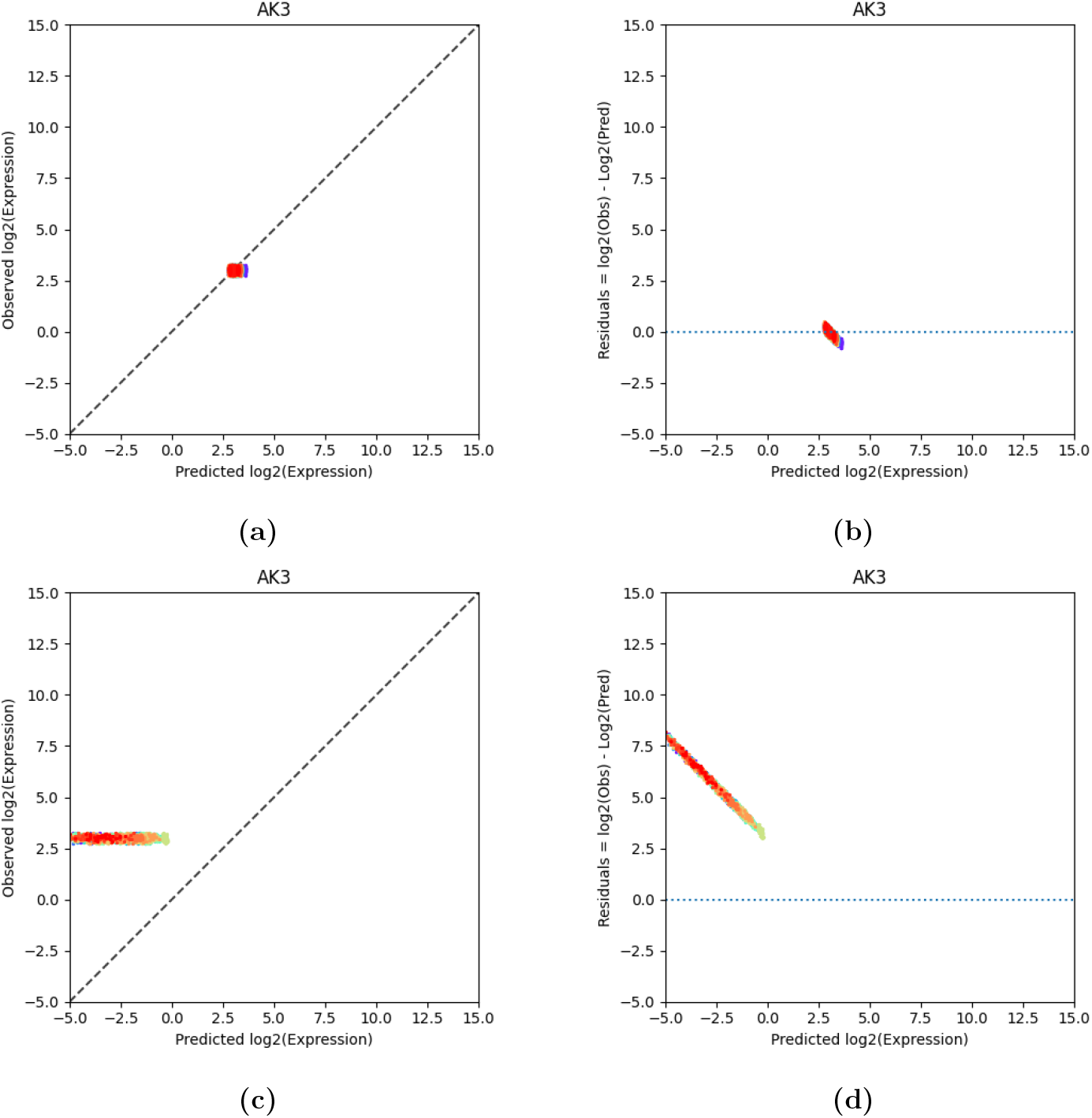
(a) Predicted versus observed expression values and (b) residuals for the test samples for all 100 shuffles for AK3 using fitted parameters. (c) Predicted versus observed expression values and (d) residuals for the test samples for all 100 shuffles for AK3 using random parameters. Each shuffle is colored.

The estimated fit of the model to training data improved over iterations of the procedure. However, the means and standard deviations from the equilibrium distributions do not converge as quickly as we would like (data not shown). This slow convergence, and the necessity for repeated estimations, mean that computational time is a limiting factor. Future analyses should simulate longer to identify the appropriate cut offs given the data used, and thus improve the fit of the model parameters.

While the use of a stochastic dynamical system offers distinct advantages over more statistically-driven methods, a number of limitations of the our approach warrant discussion. First, our model is based on the assumption that epigenetic modification effects the propensity of the random process of transcription factor binding and unbinding. As seen in other studies, gene expression is a complex mechanisms that involves other epigenetic (e.g., histone modifications and non-coding RNAs) and genetic (e.g., DNA sequence variations) factors and varies across tissues and with age. Next, our model assumes that DNA transcription is a comparatively fast (and so approximated as deterministic) process that depends on transcription factor binding. Finally, our model implicitly assumes that processes of transcription of DNA to RNA and translation from RNA to the functional protein products are immediate. Finally, we limit the scope of our testing to linear production of DNA transcript, depending on transcription factor binding status. Future efforts will be focused on improving the prediction accuracy, improving prediction robustness across training sets, improving computational efficiency, and evaluating across other gene regulatory networks, gene sets, and datasets.

By using a dynamical systems approach, our model generates an estimation of gene expression given DNA methylation based on the mechanistic hypothesis of differential binding affinity of a transcription factor caused by epigenetic modification. Our model provides predictions based directly on the biological hypotheses presented by the GRN thereby providing an easy to identify potential mechanistic hypotheses for their predictions (i.e., the binding of TF to specific sites). In addition to gene expression predictions, the characteristics of the dynamical systems approach offers multiple additional opportunities for future evaluation. First, the dynamical systems approach allows study of complex regulatory networks, including those which contains cycles. The GRN used for evaluation was acyclic. Next, in predicting gene expression our model also predicts gene regulatory activity in the form of the boolean variables *Bi*(*t*), which may be interpreted as the bound/unbound state of a regulatory protein at some DNA binding site. Using this information, we expect that our model will provide insight beyond gene expression prediction by identifying specific differential regulatory activity (e.g., which regulatory sites are bound and to what extent). Finally, our model can also be used to predict the effects of changes in methylation states at particular sites on gene expression levels. By perturbing one area of the network (e.g., a binding site), the effects on the rest of the network can be predicted (e.g., differences in regulatory activity due to epigenetic characteristics of tumor versus normal tissues).

In conclusion, we developed a dynamical system model for predicting gene expression using a gene regulatory network and epigenome data. To our knowledge, this is the first study to develop and evaluate a stochastic dynamical systems model predicting gene expression levels from epigenetic data for a given GRN. Using our model, we were able to accurately predict gene expression levels from methylation data and outperformed linear regression models. Future applications of our method will include an evaluation of the additional opportunities offered by the characteristics of a dynamical systems approach including: (1) acyclic GRNs, (2) gene regulatory activity (i.e., binding), and (3) prediction of network perturbations.

## Supporting information

**S1 File. Further details of the method.** Available on Zenodo https://doi.org/10.5281/zenodo.4441111.

**S2 Code Repository. Method source code & sample data.** Available at GitHub https://github.com/kordk/stoch_epi_lib with demonstration data available from Synapse https://www.synapse.org/#!Synapse:syn22255244/files.

## Acknowledgments

This project was initially conceived as an interdisciplinary project as part of the “Short Course in Systems Biology - a foundation for interdisciplinary careers” at the Center for Complex Biological Systems at the University of California Irvine held Jan. 21 - Feb. 8, 2019 in Irvine, CA (NIH GM126365). This work was supported by the National Cancer Institute at the National Institute of Health under Grant CA134900.

## Notes

https://doi.org/10.5281/zenodo.3970970

https://github.com/kordk/stoch_epi_lib

### Competing Interest Statement

The authors have declared no competing interest.

