## Supplemental File 1 for "A dynamical system model for predicting gene expression from the epigenome"

### 1 Methods

#### 1.1 Model Assumptions

Our model assumes fundamentally that transcription of DNA is (relatively) fast and done at a linear rate determined by the bound or unbound state of transcription factor binding sites. We assume that binding and unbinding of transcription factors is a (relatively) slow and stochastic process, with propensity proportional to availability of transcription factor. Our model is built on the hypothesis that the propensity of transcription factor binding is influenced non-linearly by epigenetic modification of the binding site. This assumption appears as the rational function

$$\frac{\mu_i}{\mu_i + (\alpha_i)^{\nu_i}}$$

in eq. (1).

We assume that "background" transcription (i.e. transcription occurring when no transcription factor is bound) is only enough to allow down regulation to exist. Therefore, we do not estimate the parameter $\gamma$ in eq. (2), instead assuming it to be the minimum value so that

$$g_j(t) = 0 \Rightarrow \frac{d}{dt} g_j \geq 0$$

In model training and testing, genes with missing expression data are assigned 'zero' expression values.

#### 1.2 Model Equations

We model gene regulation using a piecewise-deterministic Markov process (PDMP) as introduced in Davis 1984 [26] (see also [27, 28]) given by the equations:

$$B_i(t) = B_i(0) + Y_1^i\left(\int_0^t (1 - B_i(\tau))\lambda_i \frac{\mu_i}{\mu_i + (\alpha_i)^{\nu_i}}(\boldsymbol{\kappa}_i \cdot \boldsymbol{g})d\tau\right) - Y_2^i\left(\int_0^t \hat{\lambda}_i B_i(\tau)d\tau\right) \quad (1)$$

and

$$\frac{d}{dt} g_j = \gamma_j + (\boldsymbol{\phi}_j \cdot \boldsymbol{B}) - d_j g_j \quad (2)$$

where $B_i(t) \in \{0, 1\}$, is a boolean random variable representing the bound/unbound state of a binding site region of DNA and $g_i$ is the transcript amount the genes modeled. Equation (1) is given as the sum of two Poisson jump processes $Y_1^i(h_1^i(t))$ and $Y_2(h_2^i(t))$ which take values in $\mathbb{Z}_{\geq 0}$, and are peicewise constant between randomly spaced discrete time points ("jumps") [29]. The process $Y_1$ can be interpreted as the number of times a transcription factor has bound the site $i$, and the process $Y_2$ can be interpreted as the number of times a transcription factor bound to the site has dissociated. The propensities $h_1^i(t)$ and $h_2^i(t)$ have the property that if $B_i(t) = 1$, then $h_1^i(t) = 0$ and the process $Y_1^i(h_1^i(t))$ will not increase. Similarly, if $B_i(t) = 0$, $Y_2^i(h_2^i(t))$ will not increase. The result is that

$$Y_1^i(h_1^i(t)) - Y_2(h_2^i(t)) \in \{0, 1\}$$

for all $t$ provided $B_i(0) \in \{0, 1\}$, and consequently $B_i(t) \in \{0, 1\}$ for all $t$.

The propensities $h_1^i(t)$ and $h_2^i(t)$ are taken to be linear functions of the available transcription factors, which is assumed to be the same as the transcript variables $g_j$. We take the values $\kappa_{ij} \in \{0, 1\}$; these parameters along with the set of $\phi_{ji}$ represent the structure of the underlying gene regulatory network.

We include the term

$$\frac{\mu_i}{\mu_i + (\alpha_i)^{\nu_i}} \tag{3}$$

to represent the impact of epigenetic modification on the binding propensity of transcription factors. In this term, $\alpha_i$ is the measured epigenetic modification to a transcription factor binding site (e.g. percentage of methylated bases). Equation (3) is a sigmoidal function which is either strictly increasing or strictly decreasing depending on the sign of $\nu_i$. If $\nu_i > 0$, then this term decreases, implying that epigenetic modification decreases transcription factor binding. Conversely, if $\nu_i < 0$, the model implies that epigenetic modification increases transcription factor binding.

Finally, we use a linear ODE for the value of the transcripts $g_j$. We take $\phi_{ji} \in \{-1, 0, 1\}$ based on the structure of the underlying gene regulatory network. We include baseline transcription $\gamma_j$ and decay $d_j$. Because we use a linear ODE in eq. (2), we can solve exactly between jumps of $\boldsymbol{B}$. If the vector $\boldsymbol{B}$ changes at times $t^k$, then for $t \in [t^k, t^{k+1})$, we can integrate to arrive at the following formula:

$$g_j(t) = e^{-d_j(t-t^k)} \left( g_j(t^k) - \frac{\gamma_j + \sum_i \phi_{ji} B_i(t^k)}{d_j} \right) + \frac{\gamma_j + \sum_i \phi_{ji} B_i(t^k)}{d_j} \tag{4}$$

##### 1.3 Master Equation and Equilibrium Distribution

It is common practice in the study of reaction networks modeled as stochastic jump processes to represent the process using so called "chemical master equation" [**?** 31], which is the Kolmogorov forward equation for the jump process:

$$P'(\boldsymbol{x}, t) = P(\boldsymbol{x}, t)A \tag{5}$$

where $P(\boldsymbol{x}, t)$ is a row vector such that $P_j$ represents the probability that the system $\boldsymbol{x}(t)$ is in state $j$ at time $t$, and $A$ is the *generator matrix*, representing state transitions in infinitesimal time.

Note that taking $P$ to be a row vector in the notation above implies that the state space of the system is at most countably infinite. This is not the case of the PDMP, which takes values in $E = \{0, 1\}^{|\boldsymbol{B}|} \times \mathbb{R}_{\geq 0}^{|\boldsymbol{g}|}$. However, the generator for a PDMP can be defined (see Azaïs 2014 [32] for details). We can define a density $P(B^i, \boldsymbol{g}(t), t) = P^i(\boldsymbol{g})$, $i = 1, ..., |\boldsymbol{B}|$ for each possible state $\boldsymbol{B}^i$ of $\boldsymbol{B}$ such that $\sum_{i=1}^{|\boldsymbol{B}|} P^i(\boldsymbol{g}) = P(\boldsymbol{g})$ is the probability distribution for the vector $\boldsymbol{g}$, and each $P_i$ satisfies

$$\frac{dP^i(\boldsymbol{g}, t)}{dt} = \left( \frac{d\boldsymbol{g}}{dt} \right) \cdot \nabla_{\boldsymbol{g}} P^i(\boldsymbol{g}, t) + \sum_{ij} Q_{ij}(\boldsymbol{g})(P^j(\boldsymbol{g}, t) - P^i(\boldsymbol{g}, t)) \tag{6}$$

where $Q_{ij}$ represents the jump rate from state $i$ to state $j$. Rewriting this using eqs. (1) and (2), we have

$$\frac{dP^i(\boldsymbol{g},t)}{dt} = \sum_{j=1}^{|\boldsymbol{g}|}\left(\gamma_j + (\phi_j \cdot \boldsymbol{B}^i) - d_j g_j\right)\frac{\partial P^i(\boldsymbol{g},t)}{\partial g_j}$$
$$+ \sum_{(j:\|\boldsymbol{B}^j-\boldsymbol{B}^i\|_1=1)}\sum_{k=1}^{|\boldsymbol{B}|}\left[B_k^i(1-B_k^j)\lambda_k \frac{\mu_k}{\mu_k+(\alpha_k)^{\nu_k}}(\boldsymbol{\kappa}_k\cdot\boldsymbol{g}) + \hat{\lambda}_k B_k^j(1-B_k^i)\right]P^j(\boldsymbol{g},t)$$
$$- \sum_{k=1}^{|\boldsymbol{B}|}\left[(1-B_k^i)\lambda_k \frac{\mu_k}{\mu_k+(\alpha_k)^{\nu_k}}(\boldsymbol{\kappa}_k\cdot\boldsymbol{g}) + \hat{\lambda}_k B_k^i\right]P^i(\boldsymbol{g},t) \quad (7)$$

and note that in the second term, only one of the terms in the sum

$$\sum_{k=1}^{|\boldsymbol{B}|}\left[B_k^i(1-B_k^j)\lambda_k \frac{\mu_k}{\mu_k+(\alpha_k)^{\nu_k}}(\boldsymbol{\kappa}_k\cdot\boldsymbol{g}) + \hat{\lambda}_k B_k^j(1-B_k^i)\right]$$

is non-zero.

#### 1.4 Model Sampling & Equilibrium Distribution Estimation

In a discrete space, continuous time Markov process, we can estimate the equilibrium distribution as

$$f(X) = \frac{1}{T}\int_{t_0}^{T} \mathbf{1}_X(y(t))dt$$

where $y(t)$ is a sample path and $\mathbf{1}_A$ is the indicator function on the state $X$. For our continuous space PCMP we can estimate the equilibrium distribution using kernel density estimation (KDE) with a Gaussian kernel as follows:

$$f(x) = \frac{1}{T}\int_{t_0}^{T}\frac{1}{(\sqrt{2\pi}h)^d}e^{-(\boldsymbol{x}-\boldsymbol{y}(t))^T R(\boldsymbol{x}-\boldsymbol{y}(t))/2h^2}dt$$

where $R$ is some bandwidth matrix, we use the identity.

We would like to compute the marginal distributions on the various gene variables, which we can estimate with the 1 dimensional kernel

$$f_i(x) = \frac{1}{T}\int_{t_0}^{T}\frac{1}{\sqrt{2\pi}h}e^{-(x-g_i(t))^2/2h^2}dt$$

If we compute a realization, we have the jump times such that $[t_0,t_1)\sqcup[t_1,t_2)\sqcup\cdots\sqcup[t_{n-1},T) = [t_0,T)$ and so can compute

$$f_i(x) = \sum_{k=0}^{n-1}\int_{t_k}^{t_{k+1}}\frac{1}{\sqrt{2\pi}h}e^{-(x-g_i(t))^2/2h^2}dt$$

because we have $g(t)$ between jumps. We must therefore compute

$$f_i(x) = \sum_{k=0}^{n-1}\int_{t_k}^{t_{k+1}}\frac{1}{\sqrt{2\pi}h}\exp\left(-\frac{\left(x-\left[e^{-d_i(t-t_k)}(g_i(t_k)-S_i^k)+S_i^k\right]\right)^2}{2h^2}\right)dt \quad (8)$$

where

$$S_i^k = \frac{\gamma_i + \sum_j \phi_{ij}B_j}{d_i}.$$

In fig. 1, we show schematically how $f_i(x)$ is estimated from a realization of the process $\boldsymbol{B}(t), \boldsymbol{g}(t)$.

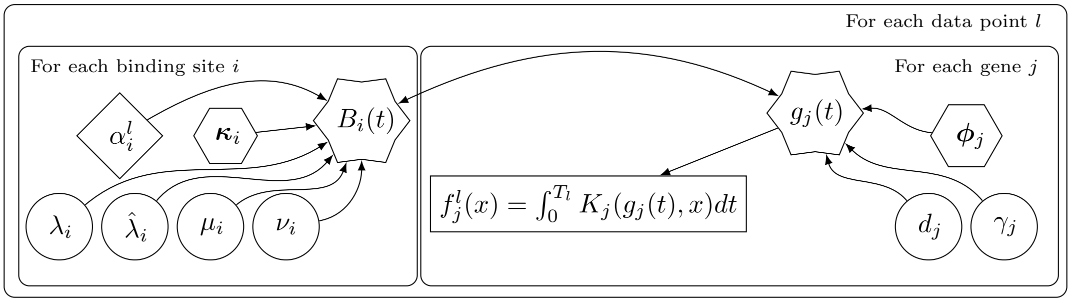


***Figure 1:*** *Plate diagram of the process to estimate the marginal PDF $f_j(x)$ of each gene's transcript level according to our model. Parameters in diamonds are read from data, parameters in hexagons are determined by the structure of the network, parameters in circles must be fit to the model by maximizing likelihood over a training data set, and parameters in stars are the state variables of the dynamical model. Notice that the dynamical model implies that the state variables depend on each other, meaning this network of dependence is* not acyclic. *The kernal $K_j(x, y)$ used to estimate likelihood is Gaussian kernel in only component $j$ (i.e., it is the Guassian kernel after orthogonal projection onto dimension $j$).*

#### 1.5 Model Parameter Estimation

##### 1.5.1 Gradient Estimation

In order to estimate $\nabla \hat{L}_{D,\boldsymbol{\omega}}$ for use in optimization, we can use the generator of the system. First, however, we redefine the likelihood estimate according to states $\boldsymbol{B}^i$ of $\boldsymbol{B}$ as

$$L^i_{\overline{\boldsymbol{g}},\boldsymbol{\alpha},\boldsymbol{\omega}}(\boldsymbol{\theta}) = \frac{1}{T}\sum_{k=0}^{n-1}\int_{t_k}^{t_{k+1}} \frac{1}{(2\pi)^{d/2}h^d} e^{-\frac{1}{2}\|(\boldsymbol{g}_{\boldsymbol{\theta},\boldsymbol{\alpha}}(t) - \overline{\boldsymbol{g}})\|^2} \delta_{\boldsymbol{B}(t),\boldsymbol{B}^i} dt \tag{9}$$

where $\delta$ is the Kronecker delta. Then,

$$L_{\overline{\boldsymbol{g}},\boldsymbol{\alpha},\boldsymbol{\omega}}(\boldsymbol{\theta}) = \sum_i L^i_{\overline{\boldsymbol{g}},\boldsymbol{\alpha},\boldsymbol{\omega}}(\boldsymbol{\theta}).$$

We will compute $\frac{\partial \hat{L}_{D,\boldsymbol{\omega}}}{\partial \theta_i}$ by first noting

$$\frac{\partial \hat{L}_{D,\boldsymbol{\omega}}}{\partial \theta_i} = -\sum_{l=1}^{n} \frac{1}{L_{\overline{\boldsymbol{g}}^l,\boldsymbol{\alpha}^l,\boldsymbol{\omega}}} \sum_i \frac{\partial}{\partial \theta_j} L^i_{\overline{\boldsymbol{g}}^l,\boldsymbol{\alpha}^l,\boldsymbol{\omega}} \tag{10}$$

and estimating $\frac{\partial}{\partial \theta_j} L^i_{\overline{\boldsymbol{g}}^l,\boldsymbol{\alpha}^l}$. For the remainder of this section, we drop the subscript $\overline{\boldsymbol{g}}_l, \boldsymbol{\alpha}^l$ and assume we are fitting to a single data point for less notational clutter. To estimated the gradient, we use the generator of the system, and assume that

$$\frac{d}{dt}L = 0$$

because we are closely approximating an equilibrium distribution. In fact, an equilibrium distribution implies the stronger condition that for each $i$, $\frac{dL^i}{dt} = 0$.

Then we have

$$\begin{aligned} 0 = &\sum_{n=1}^{|\boldsymbol{g}|} \left(\gamma_n + (\phi_n \cdot \boldsymbol{B}^i) - d_n g_n\right) \frac{\partial L^i}{\partial g_n} \\ &+ \sum_{(j:\|\boldsymbol{B}^j - \boldsymbol{B}^i\|_1 = 1)} \sum_{k=1}^{|\boldsymbol{B}|} \left[ B_k^i (1 - B_k^j) \lambda_k \frac{\mu_k}{\mu_k + (\alpha_k)^{\nu_k}} (\boldsymbol{\kappa}_k \cdot \boldsymbol{g}) + \hat{\lambda}_k B_k^j (1 - B_k^i) \right] L^j \\ &- \sum_{k=1}^{|\boldsymbol{B}|} \left[ (1 - B_k^i) \lambda_k \frac{\mu_k}{\mu_k + (\alpha_k)^{\nu_k}} (\boldsymbol{\kappa}_k \cdot \boldsymbol{g}) + \hat{\lambda}_k B_k^i \right] L^i. \end{aligned} \tag{11}$$

Differentiating with respect to some parameter $\theta_j$ we have

$$0 = \sum_{n=1}^{|\boldsymbol{g}|} \left[ \left(\gamma_n + (\phi_n \cdot \boldsymbol{B}^i) - d_n g_n\right) \frac{\partial}{\partial g_n} \left( \frac{\partial L^i}{\partial \theta_m} \right) + \frac{\partial}{\partial \theta_m} \left(\gamma_n + (\phi_n \cdot \boldsymbol{B}^i) - d_n g_n\right) \left( \frac{\partial L^i}{\partial g_n} \right) \right]$$
$$+ \sum_{(j : \|\boldsymbol{B}^j - \boldsymbol{B}^i\|_1 = 1)} \sum_{k=1}^{|\boldsymbol{B}|} \frac{\partial}{\partial \theta_m} \left( B_k^i (1 - B_k^j) \lambda_k \frac{\mu_k}{\mu_k + (\alpha_k)^{\nu_k}} (\boldsymbol{\kappa}_k \cdot \boldsymbol{g}) + \hat{\lambda}_k B_k^j (1 - B_k^i) \right) L^j$$
$$+ \sum_{(j : \|\boldsymbol{B}^j - \boldsymbol{B}^i\|_1 = 1)} \sum_{k=1}^{|\boldsymbol{B}|} \left( B_k^i (1 - B_k^j) \lambda_k \frac{\mu_k}{\mu_k + (\alpha_k)^{\nu_k}} (\boldsymbol{\kappa}_k \cdot \boldsymbol{g}) + \hat{\lambda}_k B_k^j (1 - B_k^i) \right) \frac{\partial}{\partial \theta_m} L^j$$
$$- \sum_{k=1}^{|\boldsymbol{B}|} \frac{\partial}{\partial \theta_m} \left[ (1 - B_k^i) \lambda_k \frac{\mu_k}{\mu_k + (\alpha_k)^{\nu_k}} (\boldsymbol{\kappa}_k \cdot \boldsymbol{g}) + \hat{\lambda}_k B_k^i \right] L^i$$
$$- \sum_{k=1}^{|\boldsymbol{B}|} \left[ (1 - B_k^i) \lambda_k \frac{\mu_k}{\mu_k + (\alpha_k)^{\nu_k}} (\boldsymbol{\kappa}_k \cdot \boldsymbol{g}) + \hat{\lambda}_k B_k^i \right] \frac{\partial}{\partial \theta_m} L^i. \quad (12)$$

From a sample path, we may estimate every quantity in this equation other than

$$\frac{\partial}{\partial \theta_j} L^i \ \& \ \frac{\partial}{\partial g_n} \left( \frac{\partial L^i}{\partial \theta_j} \right).$$

If $\theta_j$ is a site parameter we make the simplifying assumption that

$$\frac{\partial}{\partial g_n} \left( \frac{\partial L^i}{\partial \theta_j} \right) = 0.$$

We are then left with a linear system of equations whose solution is $\frac{\partial}{\partial \theta_j} L^i$, $i = 1, ..., 2^{|\boldsymbol{B}|}$. We note that make the above simplification to reduce the computational cost of the gradient estimate. For a more accurate estimate, one could search for an appropriate boundary condition and numerically compute the solution to the full partial differential equation. However, we see good improvement using gradient descent with the less costly gradient estimate.

Now, it will be convenient to introduce the notation $\boldsymbol{B}^{i\Delta m}$ to indicate the state of the vector $\boldsymbol{B}$ which differs from $\boldsymbol{B}^i$ only in component $m$. Note that if $L^j = L^{i\Delta m}$ then $L^i = L^{j\Delta m}$. Then, if $\theta_m \in \{\lambda_m\} \cup \{\hat{\lambda}_m\} \cup \{\mu_m\} \cup \{\nu_m\}$, then eq. (12) becomes

$$0 = \frac{\partial}{\partial \theta_m} \left( B_m^i (1 - B_m^{i\Delta m}) \lambda_m \frac{\mu_m}{\mu_m + \alpha_m^{\nu_m}} (\boldsymbol{\kappa}_m \cdot \boldsymbol{g}) + \hat{\lambda}_m B_m^{i\Delta m} (1 - B_m^i) \right) L^{i\Delta m}$$
$$+ \frac{\partial L^{i\Delta m}}{\partial \theta^m} \left( B_m^i (1 - B_m^{i\Delta m}) \lambda_m \frac{\mu_m}{\mu_m + \alpha_m^{\nu_m}} (\boldsymbol{\kappa}_m \cdot \boldsymbol{g}) + \hat{\lambda}_m B_m^{i\Delta m} (1 - B_m^i) \right)$$
$$- \frac{\partial}{\partial \theta_m} \left( (1 - B_m^i) \lambda_m \frac{\mu_m}{\mu_m + \alpha_m^{\nu_m}} (\boldsymbol{\kappa}_m \cdot \boldsymbol{g}) + \hat{\lambda}_m B_m^i \right) L^i$$
$$- \frac{\partial L^i}{\partial \theta_m} \left( (1 - B_m^i) \lambda_m \frac{\mu_m}{\mu_m + \alpha_m^{\nu_m}} (\boldsymbol{\kappa}_m \cdot \boldsymbol{g}) + \hat{\lambda}_m B_m^i \right). \quad (13)$$

Thus, in the case of site parameters, this decouples into a set of systems of two variables $L^i, L^{i\Delta m}$ that can be easily solved (all other variables in eq. (13) can be estimated from the stochastic simulation).

If $\theta_j \in \{d_n\}$ (or indeed $\theta_j$ is some other "gene" parameter in a more complicated model), then eq. (12) does not provide a way to estimate $\frac{\partial L^i}{\partial \theta_j}$. In this case, we must use a rougher estimate. We can do this by optimizing $L^i$ on a single stater $B^i$, in which case we are simply fitting parameters to an ODE model, a task for which many algorithms exist. We then estimate $\frac{\partial L}{\partial \theta_j}$ by the weighted average of these optima, weighted by the time spent in each state in a long-time realization.

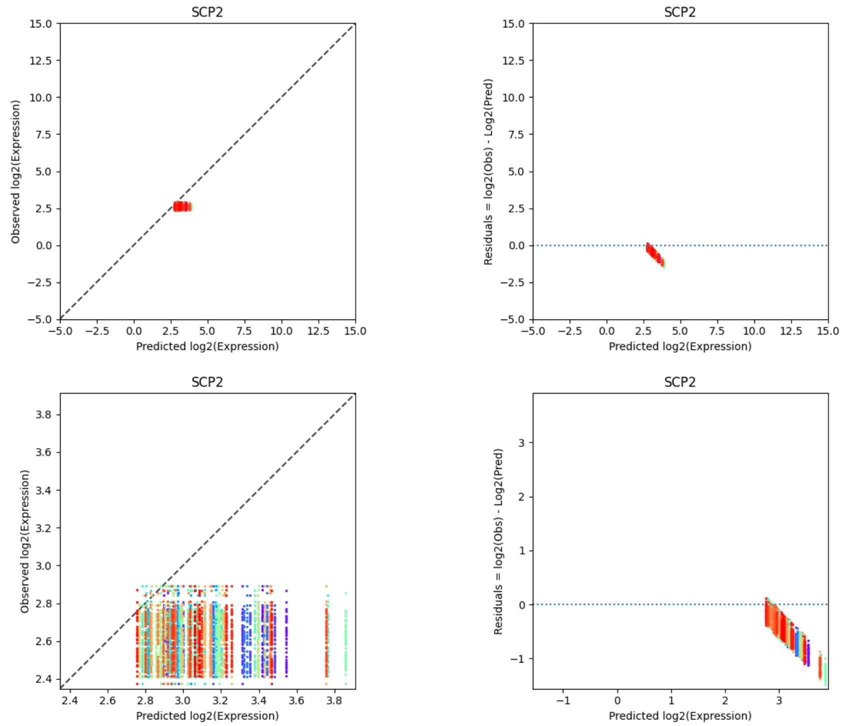


***Figure 2:*** *(left) Predicted versus observed expression values and (right) residuals for the test samples for all 100 shuffles for SCP2. Each shuffle is colored. Bottom is detailed view of top.*

#### 1.6 Additional Figures

Below, we present additional detailed results from our model evaluation.

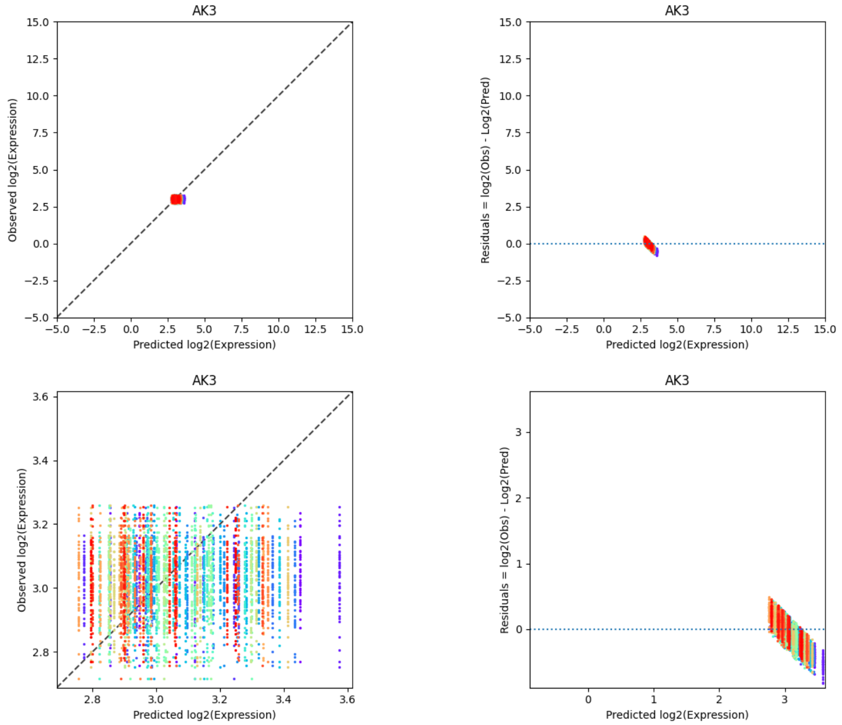


***Figure 3:*** *(left) Predicted versus observed expression values and (right) residuals for the test samples for all 100 shuffles for AK3. Each shuffle is colored. Bottom is detailed view of top.*

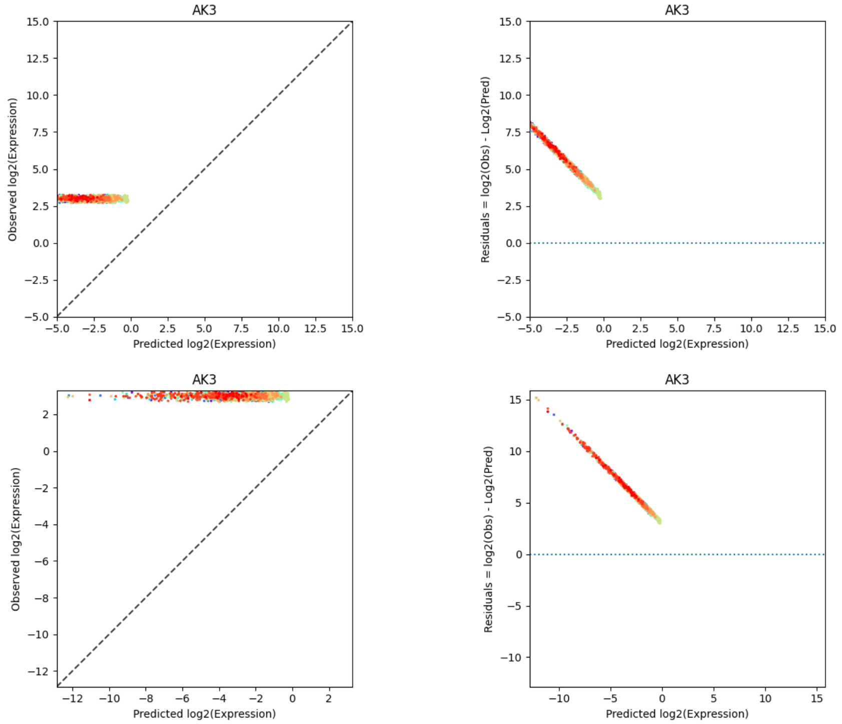


***Figure 4:*** *(Left) Predicted versus observed expression values and (Right) residuals for the test set generated from 10 sets of random parameters for all 100 shuffles for AK3. Each shuffle and parameter set is colored. Bottom is detailed view of top.*
